## Supplementary file 1 for "Membrane binding properties of the cytoskeletal protein bactofilin"

**Supplementary file 1. Diffusion constants of different BacA-mVenus variants.**

| Variant | No. of cells | No. of tracks | $D$ | $D_1$ | $D_2$ |
|---|---|---|---|---|---|
| **WT** | 130 | 2723 | 0.06 ± 0.018 | 0.02 ± 0.001 | 0.27 ± 0.001 |
| **mVenus** | 101 | 1809 | 0.29 ± 0.015 | 0.50 ± 0.002 | 1.10 ± 0.002 |
| **Δ2-8** | 120 | 2039 | 0.18 ± 0.017 | 0.24 ± 0.001 | 0.89 ± 0.001 |
| **F2Y** | 134 | 2614 | 0.13 ± 0.011 | 0.18 ± 0.002 | 0.62 ± 0.002 |
| **K4S-K7S** | 107 | 2350 | 0.08 ± 0.018 | 0.04 ± 0.001 | 0.30 ± 0.001 |
| **F2E** | 140 | 2030 | 0.17 ± 0.012 | 0.19 ± 0.001 | 0.66 ± 0.001 |
| **K4E-K7E** | 132 | 2299 | 0.25 ± 0.017 | 0.31 ± 0.002 | 0.87 ± 0.002 |
| **F2E-K4E-K7E** | 108 | 2388 | 0.16 ± 0.015 | 0.22 ± 0.001 | 0.81 ± 0.001 |
| **F130R** | 133 | 2879 | 0.22 ± 0.011 | 0.39 ± 0.001 | 0.92 ± 0.001 |
| **$^{MreB}$F130R** | 116 | 2478 | 0.26 ± 0.014 | 0.33 ± 0.002 | 0.95 ± 0.002 |
| **$^{MreB}$WT** | 123 | 2100 | 0.03 ± 0.019 | 0.01 ± 0.001 | 0.17 ± 0.001 |

$D$, MSD, average diffusion constant of all molecules ($\mu m^2 \cdot s^{-1}$)
$D_1$, diffusion constant of the slow fraction ($\mu m^2 \cdot s^{-1}$)
$D_2$, diffusion constant of the mobile fraction ($\mu m^2 \cdot s^{-1}$)
