## Supplementary file 2 for "Membrane binding properties of the cytoskeletal protein bactofilin"

**Supplementary file 2. Composition of the lipid bilayer in the MD simulations.**

| Name of lipid | Charge | Fatty acid type | Percentage share | Number count (in each layer) |
|---|---|---|---|---|
| **Phosphatidylglycerol (PG)** | | | | |
| DLPG | -1 | 12:0/12:0 | 4.68 | 6 |
| DMPG | -1 | 14:0/14:0 | 0.78 | 1 |
| DPPG | -1 | 16:0/16:0 | 1.56 | 2 |
| PYPG | -1 | 16:0/16:1 | 3.12 | 4 |
| YPPG | -1 | 16:1/16:0 | 3.12 | 4 |
| POPG | -1 | 16:0/18:1 | 1.56 | 2 |
| SOPG | -1 | 18:0/18:1 | 2.34 | 3 |
| DYPG | -1 | 16:1/16:1 | 3.91 | 5 |
| DOPG | -1 | 18:1/18:1 | 11.72 | 15 |
| **PG - total** | | | 32.81 | 42 |
| **Diacylglycerol (DAG) lipids** | | | | |
| DLGL | 0 | 12:0/12:0 | 2.34 | 3 |
| DMGL | 0 | 14:0/14:0 | 0.78 | 1 |
| DPGL | 0 | 16:0/16:0 | 2.34 | 3 |
| DSGL | 0 | 18:0/18:0 | 0.78 | 1 |
| DYGL | 0 | 16:1/16:1 | 3.12 | 4 |
| DOGL | 0 | 18:1/18:1 | 6.25 | 8 |
| **DAG - total** | | | 15.62 | 20 |
| **Monoglucosyldiglyceride (GLY)** | | | | |
| DAG-DL | 0 | 12:0/12:0 | 3.90 | 5 |
| DAG-DP | 0 | 16:0/16:0 | 7.81 | 10 |
| DAG-DY | 0 | 16:1/16:1 | 10.15 | 13 |
| DAG-DO | 0 | 18:1/18:1 | 23.44 | 30 |
| DAG-SO | 0 | 18:0/18:1 | 3.90 | 5 |
| **GLY - total** | | | 49.22 | 63 |
| **Phosphatidic acid (PA)** | | | | |
| DPPA | -1 | 16:0/16:0 | 0.78 | 1 |
| POPA | -1 | 16:0/18:1 | 0.78 | 1 |
| DOPA | -1 | 18:1/18:1 | 0.78 | 1 |
| **PA - total** | | | 2.34 | 3 |
| **TOTAL** | | | 100 | 128 |
