## Supplementary file 5 for "Membrane binding properties of the cytoskeletal protein bactofilin"

**Supplementary file 5. Strains used in this study.**

| Strain | Genotype | Construction | Source |
|---|---|---|---|
| ***C. crescentus*** | | | |
| CB15N | Synchronizable variant of wild-type strain CB15 | - | Evinger and Agabian, 1977 |
| JK5 | CB15N Δ*bacAB* | - | Kühn et al., 2010 |
| JK81 | CB15N *creS*::Tn5 | - | Laboratory stock |
| JK136 | CB15N Δ*pbpC xylX*::$P_{xyl}$-*bacA-venus* | - | Kühn et al., 2010 |
| JK281 | Δ*bacAB* Δ*pbpC* | | Kühn et al., 2010 |
| MT256 | CB15N *xylX*::$P_{xyl}$-*bacA-venus* | - | Laboratory stock |
| MT304 | Δ*pbpC* | - | Kühn et al., 2010 |
| LY70 | Δ*bacA* Δ*pbpC* | In-frame deletion of *bacA* in MT304 using pMT813 | This work |
| LY71 | Δ*bacB* Δ*pbpC* | In-frame deletion of *bacB* in MT304 using pMT815 | This work |
| LY72 | Δ*bacA* Δ*pbpC xylX*::$P_{xyl}$-*mVenus-pbpC* | Integration of pLY073 in LY70 | This work |
| LY75 | Δ*bacB* Δ*pbpC xylX*::$P_{xyl}$-*mVenus-pbpC* | Integration of pLY073 in LY71 | This work |
| LY76 | Δ*bacB* Δ*pbpC xylX*::$P_{xyl}$-*mVenus-pbpC*$_{Δ2-13}$ | Integration of pLY074 in LY71 | This work |
| LY77 | Δ*bacB* Δ*pbpC xylX*::$P_{xyl}$-*mvenus-pbpC*$_{1-13}$-*dipM*$_{224-296}$-*pbpC*$_{84-733}$ | Integration of pLY075 in LY71 | This work |
| LY84 | Δ*bacAB xylX*::$P_{xyl}$-*bacA*$_{Δ2-8}$-*mVenus* | Integration of pLY076 in JK5 | This work |
| LY88 | Δ*bacAB xylX*::$P_{xyl}$-*bacA*$_{K4S}$-*mVenus* | Integration of pLY087 in JK5 | This work |
| LY89 | Δ*bacAB xylX*::$P_{xyl}$-*bacA*$_{K4S/K7S}$-*mVenus* | Integration of pLY088 in JK5 | This work |
| LY90 | Δ*bacAB xylX*::$P_{xyl}$-*bacA-mVenus* | Integration of pLY086 in JK5 | This work |
| LY91 | Δ*bacAB xylX*::$P_{xyl}$-*bacA*$_{A6S}$-*mVenus* | Integration of pLY101 in JK5 | This work |
| LY92 | Δ*bacAB xylX*::$P_{xyl}$-*bacA*$_{K7S}$-*mVenus* | Integration of pLY102 in JK5 | This work |
| LY95 | Δ*bacAB xylX*::$P_{xyl}$-*bacA*$_{S3A}$-*mVenus* | Integration of pLY099 in JK5 | This work |
| LY96 | Δ*bacAB xylX*::$P_{xyl}$-*bacA*$_{Q5A}$-*mVenus* | Integration of pLY100 in JK5 | This work |
| LY97 | Δ*bacAB xylX*::$P_{xyl}$-*bacA*$_{F2Y}$-*mVenus* | Integration of pLY104 in JK5 | This work |
| LY103 | Δ*bacAB xylX*::$P_{xyl}$-2×*mreB*$_{1-11}$-*bacA*$_{Δ2-8}$-*mVenus* | Integration of pLY115 in JK5 | This work |
| LY111 | Δ*bacAB xylX*::$P_{xyl}$-*bacA*$_{F2E}$-*mVenus* | Integration of pLY131 in JK5 | This work |
| LY112 | Δ*bacAB xylX*::$P_{xyl}$-*bacA*$_{K4E/K7E}$-*mVenus* | Integration of pLY132 in JK5 | This work |
| LY113 | Δ*bacAB xylX*::$P_{xyl}$-*bacA*$_{F2E/K4E/K7E}$-*mVenus* | Integration of pLY138 in JK5 | This work |
| LY119 | Δ*bacAB xylX*:: $P_{xyl}$-*bacA*$_{F130R}$-*mVenus* | Integration of pLY154 in JK5 | This work |
| LY120 | *creS*::Tn5 *xylX*::$P_{xyl}$-*creS-mNeonGreen* | Integration of pLY149 in JK81 | This work |
| LY121 | *creS*::Tn5 *xylX*::$P_{xyl}$-*creS*$_{Δ2-27}$-*mNeonGreen* | Integration of pLY144 in JK81 | This work |
| LY122 | *creS*::Tn5 *xylX*::$P_{xyl}$-*bacA*$_{1-8}$-*creS*$_{28-457}$-*mNeonGreen* | Integration of pLY145 in JK81 | This work |
| LY123 | Δ*bacAB xylX*::$P_{xyl}$-2×$^{Ec}$*mreB*$_{1-11}$-*bacA*$_{F130R/9-161}$-*mVenus* | Integration of pLY155 into JK5 | This work |
| MAB568 | Δ*bacAB* Δ*pbpC vanA*::$P_{van}$-*pbpC*$_{1-132}$-*mCherry* | Integration of pMAB234 into JK281 | This work |
| MAB575 | Δ*bacAB* Δ*pbpC vanA*::$P_{van}$-*pbpC*$_{1-132}$-*mCherry xylX*::$P_{van}$-*bacA*$_{Δ2-8}$-*mVenus* | Integration of pLY76 into MAB568 | This work |
| MAB576 | Δ*bacAB* Δ*pbpC vanA*::$P_{van}$-*pbpC*$_{1-132}$-*mCherry xylX*::$P_{van}$-*bacA-mVenus* | Integration of pLY86 into MAB568 | This work |
| MAB577 | Δ*bacAB* Δ*pbpC vanA*::$P_{van}$-*pbpC*$_{1-132}$-*mCherry xylX*::$P_{van}$-*bacA*$_{F130R}$-*mVenus* | Integration of pLY154 into MAB568 | This work |
| ***E. coli*** | | | |
| TOP10 | F⁻ *mcrA* Δ(*mrr-hsd*RMS-*mcr*BC) Φ80*lacZ*ΔM15 Δ*lac*X74 *rec*A1 *ara*D139 Δ(*ara leu*) 7697 *gal*U *gal*K *rps*L (StrR) *end*A1 *nup*G | - | Invitrogen |
| Rosetta(DE3)pLysS | F⁻ *ompT hsdS*$_B$($r_B$- $m_B$-) *gal dcm* (DE3) pLysSRARE (Cam$^R$) | - | Merck Milipore |
