## Supplementary file 6 for "Membrane binding properties of the cytoskeletal protein bactofilin"

### Supplementary file 6. **Plasmids used in this study.**

| Plasmid | Description | Construction/Source |
|---|---|---|
| pAI039 | Integrative plasmid bearing *dipM-sfmTurquoise2*$^{ox}$ | Izquierdo-Martinez et al., 2023 |
| pBAD24 | Replicating plasmid for the expression of genes under the control of the arabinose-inducible $P_{BAD}$ promoter, $Amp^R$ | Guzman et al., 1995 |
| pET51b(+) | Plasmid for overexpressing proteins with a cleavable N-terminal Strep-II tag and a C-terminal 10xHis tag | Novagen |
| pLY009 | pTB146 bearing *bacA*$_{F130R}$ | (a) Amplification of *bacA*$_{F130R}$ from pLY154 by PCR with primers oLY001 and oLY002<br>(b) Insertion into *Sap*I/*BamH*I-treated pTB146 by Gibson assembly |
| pLY070 | pXCHYC-2 bearing *pbpC*$_{1\text{-}39nt}$-*dipM*$_{670\text{-}888nt}$-*pbpC*$_{250\text{-}396nt}$ | (a) Amplification of $P_{xyl}$-*pbpC*$_{1\text{-}39nt}$ from pMT993 by PCR with primers CS008 and oLY158<br>(b) Amplification of *dipM*$_{670\text{-}888nt}$ from pAI039 by PCR with primers oLY159 and oLY160<br>(c) Amplification of *pbpC*$_{250\text{-}396nt}$-*mCherry* from pMT993 by PCR with primers oLY161 and oLY162<br>(d) Gibson assembly of the three fragments<br>(e) Digestion of pXCHYC-2 and the assembly product with *Asc*I and *Nhe*I and subsequent ligation |
| pLY073 | pXmVENN-1 bearing *pbpC* | (a) Digestion of pMT906 with *Nhe*I and *Kpn*I<br>(b) Ligation of the released *pbpC* fragment into *Nhe*I/*Kpn*I-treated pXmVENN-1 |
| pLY074 | pXmVENN-1 bearing *pbpC*$_{\Delta 4\text{-}39nt}$ | (a) Amplification of *pbpC*$_{\Delta 4\text{-}39nt}$ from pMT906 by PCR with primers oLY169 and CC3277-rev2<br>(b) Restriction of the PCR product with *Nhe*I and *Kpn*I<br>(c) Ligation into *Nhe*I/*Kpn*I-treated pXmVENN-1 |
| pLY075 | pXmVENN-1 bearing *pbpC*$_{1\text{-}39nt}$-*dipM*$_{670\text{-}888nt}$-*pbpC*$_{250\text{-}2202nt}$ | (a) Amplification of *pbpC*$_{1\text{-}39nt}$-*dipM*$_{670\text{-}888nt}$ from pLY070 by PCR with primers CC3277-for and oLY170<br>(b) Amplification of *pbpC*$_{250\text{-}2202nt}$ from pMT906 by PCR with primers oLY171 and CC3277-rev2<br>(c) Gibson assembly of the two fragments<br>(d) Digestion of pXmVENN-1 and the assembly product with *Nhe*I and *Kpn*I and subsequent ligation |
| pLY076 | pXmVENC-2 bearing *bacA*$_{\Delta 4\text{-}32nt}$ | (a) Amplification of *bacA*$_{\Delta 4\text{-}32nt}$ from pMT812 by PCR with primers oLY172 and CC1873-rev<br>(b) Digestion with *Nde*I and *Sac*I<br>(c) Ligation into *Nde*I/*Sac*I-treated pXmVENC-2 |
| pLY086 | pXmVENC-2 bearing *bacA* | (a) Digestion of pMT812 with *Nde*I and *Sac*I<br>(b) Ligation of the released *bacA* fragment into *Nde*I/*Sac*I-treated pXmVENC-2 |
| pLY087 | pXmVENC-2 bearing *bacA*$_{K4S}$ | Site-directed mutagenesis of pLY086 by PCR with primers oLY192 and oLY193 |
| pLY088 | pXmVENC-2 bearing *bacA*$_{K4S/K7S}$ | Site-directed mutagenesis of pLY087 by PCR with primers oLY194 and oLY195 |
| pLY099 | pXmVENC-2 bearing *bacA*$_{S3A}$ | (a) Amplification of *bacA*$_{S3A}$ from a custom-synthesized gene block by PCR with primers oLY217 and oLY218<br>(b) Amplification of *bacA*$_{106\text{-}483nt}$ from pLY086 by PCR with primers oLY219 and CC1873-rev<br>(c) Gibson assembly of the two fragments<br>(d) Digestion of pXmVENC-2 and the assembly product with *Nde*I and *Sac*I and subsequent ligation |
| pLY100 | pXmVENC-2 bearing *bacA*$_{Q5A}$ | (a) Amplification of *bacA*$_{Q5A}$ from a custom-synthesized gene block by PCR with primers oLY217 and oLY218<br>(b) Amplification of *bacA*$_{106\text{-}483nt}$ from pLY086 by PCR with primers oLY219 and CC1873-rev<br>(c) Gibson assembly of the two fragments<br>(d) Digestion of pXmVENC-2 and the assembly product with *Nde*I and *Sac*I and subsequent ligation |
| pLY101 | pXmVENC-2 bearing *bacA*$_{A6S}$ | (a) Amplification of *bacA*$_{A6S}$ from a custom-synthesized gene block by PCR with primers oLY217 and oLY218<br>(b) Amplification of *bacA*$_{106\text{-}483nt}$ from pLY086 by PCR with primers oLY219 and CC1873-rev<br>(c) Gibson assembly of the two fragments<br>(d) Digestion of pXmVENC-2 and the assembly product with *Nde*I and *Sac*I and subsequent ligation |

**Supplementary file 5. Plasmids used in this study (continued).**

| Plasmid | Description | Construction/Source |
|---|---|---|
| pLY102 | pXmVENC-2 bearing $bacA_{K7S}$ | (a) Amplification of $bacA_{K7S}$ from a custom-synthesized gene block by PCR with primers oLY217 and oLY218<br>(b) Amplification of $bacA_{106\text{-}483nt}$ from pLY086 by PCR with primers oLY219 and CC1873-rev<br>(c) Gibson assembly of the two fragments<br>(d) Digestion of pXmVENC-2 and the assembly product with *Nde*I and *Sac*I and subsequent ligation |
| pLY104 | pXmVENC-2 bearing $bacA_{F2Y}$ | Site-directed mutagenesis of pLY086 by PCR with primers oLY222 and oLY223 |
| pLY107 | pET51b(+) bearing MCS-*mVenus* from pXmVENC-2 | (a) Amplification of MCS-*mVenus* from pXmVENC-2 by PCR with primers oLY227 and oLY228<br>(b) Insertion into *Nco*I/*Avr*II-treated pET51b(+) by Gibson assembly |
| pLY112 | pLY107 bearing 2x$mreB_{EC\ 1\text{-}33nt}$ | (a) Annealing of oligonucleotides oLY240 and oLY241<br>(b) Ligation into pLY107 cut with *Nde*I and *Kpn*I<br>(c) Annealing of oligonucleotides oLY242 and oLY243<br>(d) Ligation into the plasmid from step (b) cut with *Xho*I and *Eco*RI |
| pLY115 | pXmVENC-2 bearing 2x$mreB_{EC\ 1\text{-}33nt}$-$bacA_{\Delta 4\text{-}32nt}$-*mVenus* | (a) Amplification of 2x$mreB_{EC\ 1\text{-}33nt}$ from pLY112 by PCR with primers oLY249 and oLY250<br>(b) Insertion into *Nde*I-treated pLY076 by Gibson assembly |
| pLY116 | pTB146 bearing $bacA_{\Delta 4\text{-}32nt}$ | (a) Amplification of $bacA_{\Delta 4\text{-}32nt}$ from pLY076 by PCR with primers oLY251 and oLY002<br>(b) Insertion into *Sap*I/*BamH*I-treated pTB146 by Gibson assembly |
| pLY117 | pTB146 bearing $bacA_{F2Y}$ | (a) Amplification of $bacA_{F2Y}$ from pLY104 by PCR with primers oLY252 and oLY002<br>(b) Insertion into *Sap*I/*BamH*I-treated pTB146 by Gibson assembly |
| pLY118 | pTB146 bearing $bacA_{K4SK7S}$ | Amplification of $bacA_{K4SK7S}$ from pLY088 by PCR with primers oLY253 and oLY002.<br>(b) Insertion into *Sap*I/*BamH*I-treated pTB146 by Gibson assembly |
| pLY119 | pTB146 bearing *bacA* | (a) Amplification of *bacA* from pMT812 by PCR with primers oLY001 and oLY002<br>(b) Insertion into *Sap*I/*BamH*I-treated pTB146 by Gibson assembly |
| pLY131 | pXmVENC-2 bearing $bacA_{F2E}$ | Site-directed mutagenesis of pLY086 by PCR with primers oLY269 and oLY270 |
| pLY132 | pXmVENC-2 bearing $bacA_{K4E/K7E}$ | (a) Site-directed mutagenesis of in pLY086 by PCR with primers oLY277 and oLY278<br>(b) Site-directed mutagenesis of the resulting plasmid by PCR with primers oLY279 and oLY280 |
| pLY133 | pTB146 bearing $bacA_{F2E}$ | (a) Amplification of $bacA_{F2E}$ from pLY131 by PCR with primers oLY273 and oLY002<br>(b) Insertion into *Sap*I/*BamH*I-treated pTB146 by Gibson assembly. |
| pLY134 | pTB146 bearing $bacA_{xE}$ | (a) Amplification of $bacA_{xE}$ from pLY131 by PCR with primers oLY274 and oLY002<br>(b) Insertion into *Sap*I/*BamH*I-treated pTB146 by Gibson assembly |
| pLY135 | pTB146 bearing $bacA_{\Delta M}$ | (a) Amplification of $bacA_{\Delta M}$ from pLY086 by PCR with primers oLY275 and oLY002.<br>(b) Insertion into *Sap*I/*BamH*I-treated pTB146 by Gibson assembly |
| pLY136 | pTB146 bearing $bacA_{K4E/K7E}$ | (a) Amplification of $bacA_{K4EK7E}$ from pLY132 by PCR with primers oLY276 and oLY002<br>(b) Insertion into *Sap*I/*BamH*I-treated pTB146 by Gibson assembly |
| pLY138 | pXmVENC-2 bearing $bacA_{F2E/K4E/K7E}$ | (a) Site-directed mutagenesis of pLY131 by PCR with primers oLY281 and oLY278<br>(b) Site-directed mutagenesis of the resulting plasmid by PCR with primers oLY279 and oLY282 |
| pLY139 | pTB146 bearing $bacA_{F2E/K4E/K7E}$ | (a) Amplification of $bacA_{F2EK4EK7E}$ from pLY138 by PCR with primers oLY283 and oLY002<br>(b) Insertion into *Sap*I/*BamH*I-treated pTB146 by Gibson assembly |
| pLY144 | pXmNeonGreenC-4 bearing $creS_{\Delta 1\text{-}81nt}$ | Deletion of nt 1-81 of *creS* in pLY149 by inverse PCR with primers oLY287 and oLY288 |
| pLY145 | pXmNeonGreenC-4 bearing $bacA_{1\text{-}24nt}$-$creS_{82\text{-}1371nt}$ | Replacement of the first 81 nucleotides of *creS* with $bacA_{1\text{-}24nt}$ in pLY149 by inverse PCR with primers oLY289 and oLY290 |
| pLY149 | pXmNeonGreenC-4 bearing *creS* | (a) Amplification of *creS* from genomic DNA of CB15N by PCR with primers creS-F and oLY301<br>(b) restriction with *Nde*I and *Kpn*I<br>(c) Ligation with pXmNeonGreenC-4 cut with *Nde*I and *Kpn*I |
| pLY154 | pXmVENC-2 bearing $bacA_{F130R}$ | Site-directed mutagenesis of pLY086 by PCR with primers oLY004 and oLY005 |
| pLY155 | pXmVENC-2 bearing 2x$mreB_{Ec1\text{-}33nt}$-$bacA_{\ F130R/31\text{-}483nt}$ | Site-directed mutagenesis of pLY115 by PCR with primers oLY004 and oLY005 |

**Supplementary file 4. Plasmids used in this study (continued).**

| Plasmid | Description | Construction/Source |
|---|---|---|
| pMAB234 | pVGFPC-4 bearing pbpC$_{11\text{-}396nt}$-*mCherry* | a) Digestion of pMT993 with *Nde*I and *Nhe*I to isolate a fragment containing *pbpC*$_{11\text{-}396nt}$-*mCherry*<br>b) Ligation into pVGFPC-4 cut with *Nde*I and *Nhe*I |
| pMT812 | pXVENC-2 bearing *bacA* | Kühn et al., 2010 |
| pMT813 | pNPTS138 derivative used to generate an in-frame deletion in *bacA* | Kühn et al., 2010 |
| pMT815 | pNPTS138 derivative used to generate an in-frame deletion in *bacB* | Kühn et al., 2010 |
| pMT906 | pXVENN-1 bearing *pbpC* | Kühn et al., 2010 |
| pMT993 | pXCHYC-2 bearing *pbpC*$_{1\text{-}396nt}$ | Kühn et al., 2010 |
| pTB146 | Plasmid for overexpression of protein with N-terminal $His_6$-SUMO fusion, $Amp^R$ | Bendezu et al., 2009 |
| pVGFPC-4 | Integrative vector for the production of fusion proteins carrying a C-terminal eGFP tag under the control of van, $Gent^R$ | Thanbichler et al., 2007 |
| pXCHYC-2 | Integrative vector for the production of fusion proteins carrying a C-terminal mCherry tag under the control of $P_{xyl}$, $Kan^R$ | Thanbichler et al., 2007 |
| pXmVENC-2 | Integration plasmid for the production of fusion proteins carrying a C-terminal mVenus tag under the control of $P_{xyl}$, $Kan^R$ | (a) Site-directed mutagenesis of *venus* by inverse PCR using pXVENC-2 as template and primers venus-mut-for/-rev<br>(b) amplification of *venus*(A207K) from the mutagenized vector by PCR using primers Pxyl-GA-for and venus-GA-r2<br>(c) insertion of the PCR product into NdeI/NheI-treated pXVENC-2 by Gibson Assembly |
| pXmVENN-1 | Integration plasmid for the production of fusion proteins carrying an N-terminal mVenus tag under the control of $P_{xyl}$, $Strep/Spec^R$ | (a) Amplification of *mVenus* from pXmVENC-2 by PCR with primers oLY167 and oLY168<br>(b) Digestions of pXVENN-1 and the PCR product with *Nde*I and *Bsr*GI and subsequent ligation |
| pXmNeonGreenC-4 | Integration plasmid for the production of fusion proteins carrying a C-terminal mNenoGreen tag under the control of $P_{xyl}$, $Gent^R$ | (a) Amplification of *mNeonGreen* from pmNeonGreen-N1 (Allele Biotechnology) by PCR with primers oLY299 and oLY300<br>(b) Digestion of pXGFPC-4 and the PCR product with AgeI and NheI and subsequent ligation |
