## Supplementary file 7 for "Membrane binding properties of the cytoskeletal protein bactofilin"

**Supplementary file 7. Oligonucleotides used in this study.** Restriction sites are underlined.

| Oligonucleotide | Sequence (5' → 3') |
|---|---|
| CC1873-rev | TAGAGCTCCGCCGGCGCTCTTGGCGATCGCCAGA |
| CC3277-for | TTGGTACCATGAACGACTGGACGCTGCCGCCCTA |
| CC3277-rev2 | TATAGCTAGCCTAGTAGGGCAGGTTGTCCGGGGGCGG |
| cres-F | CTCGAGTTTTGGGGAGACGACCATATGAGACTGCTGTCGAAGAACTCGC |
| oLY001 | CTCACAGAGAACAGATTGGTGGTATGTTCAGCAAGCAAGCTAAATCG |
| oLY002 | GCTTTGTTAGCAGCCGGATCCTTAGCCGGCGCTCTTGGCGATC |
| oLY004 | AACCGGCGCCTTCCGCCAGGGCCGCAGC |
| oLY005 | GCTGCGGCCCTGGCGGAAGGCGCCGGTT |
| oLY169 | TTGGTACCATGGGGAAGAGTCCCGGAGAGC |
| oLY170 | GTCCTCGAAGCGAATGCCCGGCGCCCGGTTCAC |
| oLY171 | AACCGGGCGCCGGGCATTC |
| oLY172 | TATCATATGAACAACAAGGCCCCGGCCC |
| oLY192 | AGCCAAGCTAAATCGAACAACAAGGCCCCGG |
| oLY193 | GCTGAACATATGGTCGTCTCCCCAAAACTCG |
| oLY194 | AGCTCGAACAACAAGGCCCCGGCC |
| oLY195 | AGCTTGGCTGCTGAACATATGGTCGTCT |
| oLY217 | ACGCTCGAGTTTTGGGGAGACGACC |
| oLY218 | CGAGGCGACCTTGGGCGGTGCGCGACGCGCGGGCT |
| oLY222 | TACAGCAAGCAAGCTAAATCGAACAACAAGGC |
| oLY223 | CATATGGTCGTCTCCCCAAAACTCGAGCG |
| oLY227 | GTTTAACTTTAAGAAGGAGATATACCATATGCCTGCAGGCGCCTTAATTAATATGC |
| oLY228 | TCAGCGGTGGCAGCAGCCTAGGTTACTTGTACAGCTCGTCCATGCCGAGAG |
| oLY240 | TATGTTGAAAAAAATTTCGTGGCATGTTTTCCAATGGTAC |
| oLY241 | CATTGGAAAACATGCCACGAAATTTTTTCAACA |
| oLY242 | TCGAGACATGTTGAAAAAAATTTCGTGGCATGTTTTCCAATTCG |
| oLY243 | AATTCGAATTGGAAAACATGCCACGAAATTTTTTCAACATGTC |
| oLY249 | TTGGGGAGACGACCATATGATGTTGAAAAAAATTTCGTGGCATGTTTTCCAATGGTAC |
| oLY250 | GGGCCTTGTTGTTCATATGGGTGGCCGACCGGTGACGC |
| oLY251 | CTCACAGAGAACAGATTGGTGGTATGAACAACAAGGCCCCGGCC |
| oLY252 | CTCACAGAGAACAGATTGGTGGTATGTACAGCAAGCAAGCTAAATCGAACAACAAGG |
| oLY253 | CTCACAGAGAACAGATTGGTGGTATGTTCAGCAGCCAAGCTAGCTCGAAC |
| oLY269 | CGACCATATGGAGAGCAAGCAAGCTAAATC |
| oLY270 | TCTCCCCAAAACTCGAGC |
| oLY273 | CTCACAGAGAACAGATTGGTGGTATGGAGAGCAAGCAAGCTAAATCGAAC |
| oLY274 | CTCACAGAGAACAGATTGGTGGTGAGAGCAAGCAAGCTAAATCGAAC |
| oLY275 | CTCACAGAGAACAGATTGGTGGTTTCAGCAAGCAAGCTAAATCGAACAACAAGG |
| oLY276 | CTCACAGAGAACAGATTGGTGGTATGTTCAGCGAGCAAGCTGAGTCG |
| oLY277 | TATGTTCAGCGAGCAAGCTAAATC |
| oLY278 | TGGTCGTCTCCCCAAAAC |
| oLY279 | CGAGCAAGCTGAGTCGAACAACAAG |
| oLY280 | CTGAACATATGGTCGTCTC |
| oLY281 | TATGGAGAGCGAGCAAGCTAAATC |
| oLY282 | CTCTCCATATGGTCGTCTC |
| oLY283 | CTCACAGAGAACAGATTGGTGGTATGGAGAGCGAGCAAGCTGAGTCG |
| oLY287 | ATGCAGCACCAGATCGAG |
| oLY288 | CATATGGTCGTCTCCCCAAA |
| oLY289 | TAAATCGAACAACATGCAGCACCAGATCGAG |
| oLY290 | GCTTGCTTGCTGAACATATGGTCGTCTCCCCAAA |
| oLY301 | ATGCGGTACCAGGCGCTCGCGGCCACGT |
| Pxyl-GA-for | GTTTTGGGGAGACGACCATATGCCTGCAGGCGCCTTAATTAATATG |
| venus-GA-r2 | CCCCGGGCTGCAGCTAGCTTACTTGTACAGCTCGTCCATGCCGAG |
| venus-mut-for | CTGAGCTACCAGTCCAAGCTGAGCAAAGACCCCC |
| venus-mut-rev | GGGGTCTTTGCTCAGCTTGGACTGGTAGCTCAG |
